# Possible Regulation of Toll-Like Receptor 4 By Lysine Acetylation Through LPCAT2 Activity in RAW264.7 Cells

**DOI:** 10.1101/2021.11.25.469959

**Authors:** Victory Ibigo Poloamina, Wondwossen Abate, Gyorgy Fejer, Simon K. Jackson

**Affiliations:** University of Plymouth, Faculty of Health, Plymouth, UK PL4 8AA; University of Exeter, College of Medicine and Health, Exeter, UK EX1 2HZ; MolEndoTech Ltd, Brixham, UK TQ5 8BA

## Abstract

Inflammation is central to several diseases. TLR4 mediates inflammatory signals, however, there are gaps in the understanding of its mechanisms. Recently, TLR4 was found to co-localise with LPCAT2, a lysophospholipid acetyltransferase. This interaction influenced TLR4 subcellular localisation through an unknown mechanism.

In this study, we have combined computational analysis, RNA interference technology, and biochemical analysis to investigate the possibility of TLR4 lysine acetylation and the influence of LPCAT2 on the detected lysine acetylation.

The results suggest for the first time that TLR4 can undergo lysine acetylation and LPCAT2 can influence TLR4 lysine acetylation. This lays a foundation for further research on the role of lysine acetylation on TLR4 and characterisation of LPCAT2 as a protein acetyltransferase.

## 2 Introduction

Inflammation is central to many diseases such as cancer, asthma, sepsis, and cardiovascular diseases [1]. It can be caused by infection from various micro-organisms, or by cell damage [2]. During bacterial infections, TLR4 plays a major role in mediating inflammatory signals after it recognises bacterial lipopolysaccharide bound to co-receptor CD14 [3,4]. Although there is scientifically established information on TLR4 mechanisms of signalling and protein-protein interactions, a lot is yet to be understood about TLR4 mechanisms. Recently, TLR4 was found to co-localise with LPCAT2; this led to a change of its subcellular localisation [5]. Nonetheless, how LPCAT2 affects the subcellular localisation of TLR4 is not known.

LPCAT2 is known as a lipid acyltransferase and acetyltransferase [6]. It is highly expressed in inflammatory cells such as peritoneal macrophages, microglia, and neutrophils [7]. In experimental allergic encephalomyelitis [8,9] and peripheral nerve injury [10–12], LPCAT2 is significantly expressed. Furthermore, LPCAT2 is suggested as a biomarker for sepsis and allergic asthma [13–15]. On the other hand, experimental conditions where LPCAT2 is silenced or inhibited, results in the resolution of inflammation via decreased production of cytokines and LPCAT2 metabolites [5,10,16]. During inflammatory conditions, LPCAT2 and TLR4 expression increases in the lipid raft domain of RAW264.7 macrophage cell line, which serves as a platform for mediating inflammatory signals [5]. Acetylation is a post-translational modification that can influence the subcellular localisation and function of a protein [17,18]. Although it commonly occurs on histones, there are several recent scientific publications that have identified acetylation on non-histone proteins [19]. Lysine acetylation can regulate protein function, interaction, and localisation [20,21]. LPCAT1 which has a very similar structure and function to LPCAT2 is known to palmitoylate histone 4, a protein [22]. This allows for the theory that LPCAT2 could carry out either of its enzymatic activities; acyltransferase or acetyltransferase on proteins such as TLR4.

Since several scientific publications suggest that LPCAT2 participates in inflammation, understanding the molecular mechanisms of LPCAT2 will contribute new knowledge on inflammation and could lead to new therapies for inflammatory disorders.

This study uses computational analysis, RNA interference technology, and biochemical analysis to detect lysine acetylation on TLR4 and the possibility of LPCAT2 influencing the detected TLR4 lysine acetylation.

## 3 Materials and Methods

Chemicals reagents used to prepare buffers and BCA Assay kit were purchased from Sigma Aldrich, UK and Fisher Scientific, UK. Buffers used include RIPA buffer, Phosphate-buffered saline (PBS), Tris-buffered saline (TBS), Blocking Buffer, Cell lysis buffer, elution buffer, SDS sample buffer, and ECL detection reagent [23]. PolyPlus INTERFERin was purchased from Source Bioscience, UK. Pre-designed siRNA, Opti-MEM, Power SYBR Green, RNa to cDNA kit, and gel casting materials were purchased from Life Technologies, UK. Antibodies, Protein A/G agarose gel beads, and protein ladders were obtained from Santa Cruz Biotechnology, UK and Cell Signalling Technologies, UK. DMEM culture medium and other cell culture materials were purchased from Lonza, UK. PCR primers were designed with Primer3 Plus Bioinformatics Softwaew and NCBI BLAST, and purchased from Eurofins Genomics.

### 3.1 Cell line and Culture

RAW264.7 cell line was obtained from the European Collection of Cell Cultures (ECACC) through Public Health England, UK. RAW264.7 macrophages were maintained in Dulbecco’s Modified Eagle Medium (DMEM) [Lonza, BE12-914F] supplemented with 10%(v/v) Foetal Bovine Serum (FBS) [Labtech.com, BS-110] and 1%(v/v) 0.2M L-Glutamine [BE17-605E], and incubated at 37ºC, 5% CO_2_. Lipopolysaccharide (*E. coli* O111:B4) [Sigma-Aldrich, L2630] was resuspended in LAL reagent water (*<*0.005EU/ml endotoxin levels) [Lonza, W50-640]. Cell culture medium was used to dilute all ligands to the needed concentration before stimulation.

### 3.2 Transfection of RAW264.7 Cells with LPCAT2 siRNA

RAW264.7 cells were cultured 24 hours before gene silencing. Then using Opti-MEM (Reduced Serum Medium) as a diluent, a transfection mixture containing 7nM of siRNA was prepared and added to cells. The cells were incubated at 37ºC, 5% CO_2_ with Opti-MEM for 24 hours for efficient gene silencing.

### 3.3 Reverse Transcription and Real-Time Quantitative PCR

The reaction mastermix contained 37%(v/v) nuclease-free, 230nM of target primers (a mixture of both forward and reverse primers), 60%(v/v) Power SYBR Green, and 3*μ*l of ≥ 100ng/*μ*l cDNA. The reaction was initiated at 95ºC for 10 minutes, then up to 40 repeated cycles of denaturing (15 seconds, 95ºC), annealing, and extension (60 seconds, 60ºC). GAPDH and ATP5B were used as an endogenous controls.

### 3.4 Immunoprecipitation

Equal amounts of whole cell lysates were pre-cleared, and incubated with target antibodies overnight at 4ºC. Then protein A/G agarose gel beads were rinsed with PBS, added to the lysate-antibody mixture, and incubated overnight at 4ºC. The mixture was centrifuged to separate the supernatant from protein A/G agarose beads. The beads were rinsed with PBS, then eluates were made with mild to harsh elution buffers [23].

### 3.5 Immunoblotting

Equal amounts of whole cell lysates and eluates were separated on pre-cast SDS-PAGE gels and blotted on to a PVDF membrane using a blot module. The blots were blocked with 0.1% bovine serum albumin in PBS-0.1% Tween 20, probed with primary and then HRP-conjugated secondary antibodies. The target proteins separated on the PVDF membrane were detected and analysed using ImageJ.

### 3.6 Computational Analysis

Computational analysis of mouse protein sequences obtained from Uniprot database was carried out using GPS-PAIL 2.0 [24], and R Programming Software (packages used were: Peptides, Biostrings, phangorn, tidyverse, ape, seqinr, rentrez, msa, and ASEB). Both gene (rentrez) and protein sequences were aligned using multiple sequence alignment (msa). The physicochemical properties of peptides were analysed using Peptides package. Phylogenetic trees were made using maximum parsimony with SeaView version 4 [25] and phangorn package. Sequence IDs: LPCAT2-Q8BYI6, NM173014; LPCAT1-Q3TFD2, NM145376; KAT2A-Q9JHD2, NM020004; KAT2B-Q9JHD1, NM020005; CREBBP-NM001025432; ELP3-NM001253812; EP300-NM177821; KAT5-NM001362372; KAT6A-NM001081149; KAT6B-NM017479; KAT8-NM026370.

### 3.7 Data Analysis

Statistical Analysis was carried out in R Statistical Programming Software and graphs were plotted using ggplot2 package. Independent experiments were repeated at least 3 times. Data represent Mean ± Standard Error of Mean unless stated otherwise. Paired T-Test with Dunnetts T3 multiple comparison test were used for statistical analysis. All statistical tests were significant at 95% confidence interval, p ≤ 0.05.

## 4 Results

### 4.1 Transfection of RAW264.7 Cells with LPCAT2 siRNA Does Not Affect the Basal Function of RAW264.7 Cells

The gene and protein expression of LPCAT2 were analysed to confirm the knockdown of LPCAT2. Figure 1A shows that LPCAT2 gene is significantly lower in cells transfected with LPCAT2 siRNA (0.25 ± 0.02, p = 0.0015). Likewise, LPCAT2 protein decreased (4-fold) in cells transfected with LPCAT2 siRNA (Figure 1B&C).

**Figure 1:**
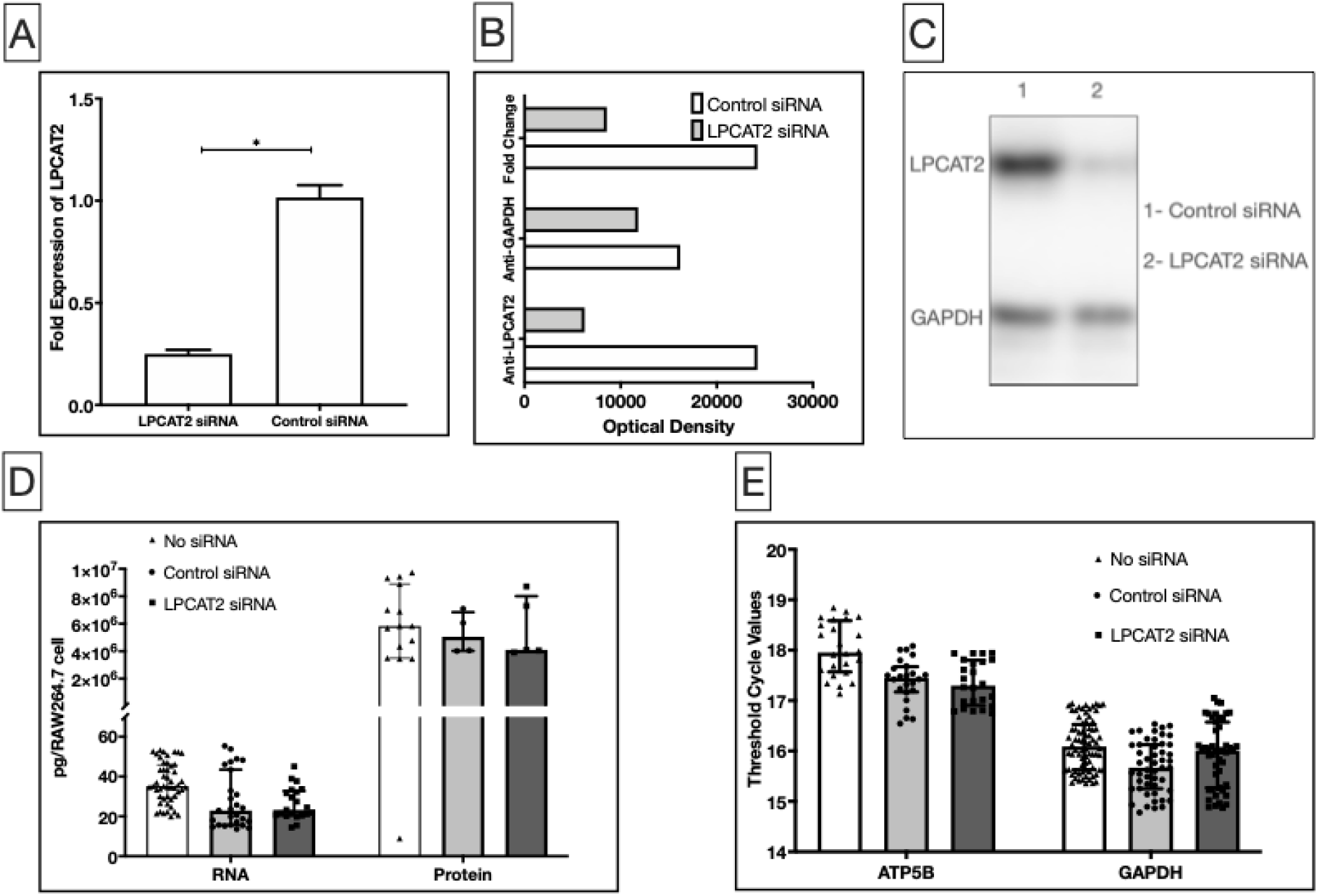
Transfection of RAW264.7 cells with LPCAT2 siRNA causes ≥70% decrease in LPCAT2 gene expression **(A)** and ≥50% decrease in LPCAT2 protein expression **(B&C)**. *p≤0.05. Fold change shows the optical density of LPCAT2 normalised to GAPDH. Data represents mean of at least 3 independent experiments (n≥3) ± standard error (A), and optical densities (B) of western blots (C). Transfection of siRNA into RAW264.7 cells does not cause a significant difference in total RNA and total protein **(D)**, and in the expression of housekeeping genes ATP5B and GAPDH **(E)** when compared to cells with no transfected siRNA. Data represents the median of at least 3 independent experiments (n≥3) ± interquartile range.

The basal function of RAW264.7 cells after transfecting the cells with LPCAT2 siRNA was analysed by measuring the total amount of RNA and Protein, and the gene expression of housekeeping genes ATP5B and GAPDH. Figure 1C shows that transfection of RAW264.7 cells with LPCAT2 siRNA does not significantly affect the total amount of RNA (23.25pg/RAW264.7 cell; 20.56pg/RAW264.7 cell to 32.78 pg/RAW264.7 cell) when compared with non-transfected cells (35.2 pg/RAW264.7 cell; 29.39pg/RAW264.7 cell to 45.84pg/RAW264.7 cell, p = 0.98) and the total amount of protein (4.09 × 10^6^pg/RAW264.7 cell; 3.98 × 10^6^ pg/RAW264.7 cell to 8.01 × 10^6^pg/RAW264.7 cell) when compared with non-transfected cells (5.82 × 10^6^pg/RAW264.7 cell; 3.5 × 10^6^pg/RAW264.7 cell to 8.89 × 10^6^pg/RAW264.7 cell, p = 0.98). Likewise, Figure 1D shows that transfection of RAW264.7 cells with LPCAT2 siRNA does not significantly affect the gene expression of housekeeping gene ATP5B (17.3; 16.9 to 17.8) when compared with non-transfected cells (17.96; 17.57 to 18.58, p = 0.95) and GAPDH (16.01; 15.27 to 16.57) when compared with non-transfected cells (16.09; 16.52 to 15.63, p = 0.95).

### 4.2 Analysis of The Relatedness of LPCAT2 to Commonly Known Lysine Acetyltransferases (KATs)

Using R Statistical Programming Software, the genetic relatedness of LPCAT2 to other KATs was analysed. As LPCAT1 is a very similar protein to LPCAT2, it was included as a positive control. Indeed, Figure 2A shows that LPCAT2 and LPCAT1 belong to the same family. Moreover, it suggests that LPCAT2 and LPCAT1 belong to the same superfamily as KAT2A and KAT2B. Therefore, the similarity of the protein sequences was analysed by aligning LPCAT2 to each protein from node 7 in Figure 2A. Figure 2B-D shows that LPCAT2 vs LPCAT1 has less than 5% gaps in alignment (B), whereas, LPCAT2 vs KAT2A (C) and LPCAT2 vs KAT2B (D) have less than or equal to 35% gaps in alignment.

**Figure 2:**
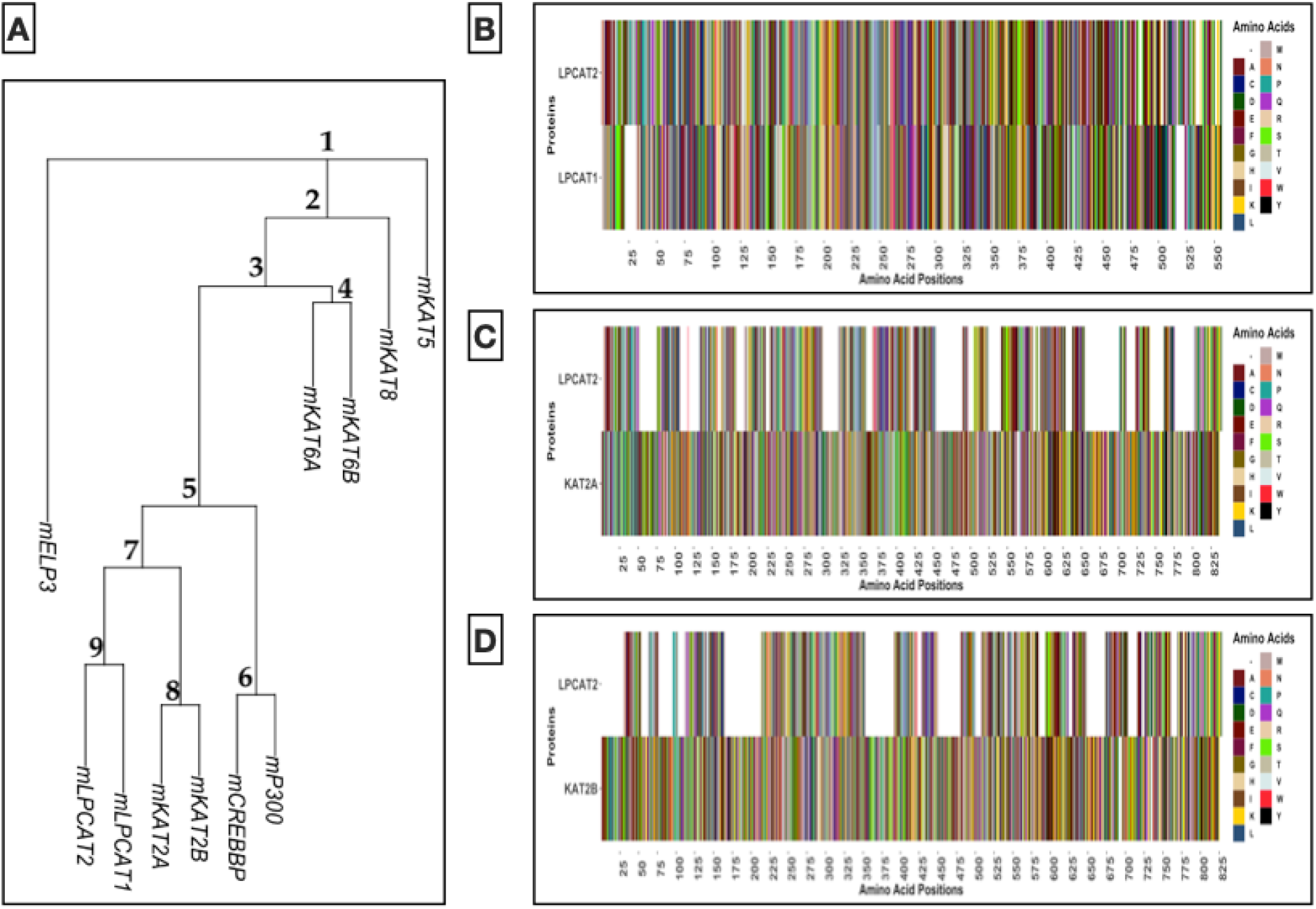
Analysis of The Relatedness of LPCAT2 To Other Lysine Acetyltransferases in Mice. **A** – Phylogenetic tree showing the degree of relatedness of LPCAT1 gene, LPCAT2 gene, and genes of other KATs. Numbers indicate node positions. **B to D** – Sequence alignment of proteins in node 7, white space indicates gaps in alignment. Letters symbolising amino acids are IUPAC standards.

### 4.3 LPCAT2 Influences Pan-Lysine Acetylation in RAW264.7 Cells

Acetylated lysine residues were detected using acetylated lysine antibodies. Figure 3 shows that stimulating RAW264.7 cells with LPS increased the density of some acetylated lysine protein bands, especially around 10kDa. The bars show fold change after normalising optical densities to acetylated alpha tubulin densities. Some protein bands (100kDa and ∼5kDa) showed ≥50% reduction after knockdown of LPCAT2 with or without LPS (Figure 3C). Although the median is very similar across all treatment, the upper quartile is lower when LPCAT2 is silenced (Figure 3B). This indicates lower band intensities of lysine acetylated proteins after LPCAT2 knockdown.

**Figure 3:**
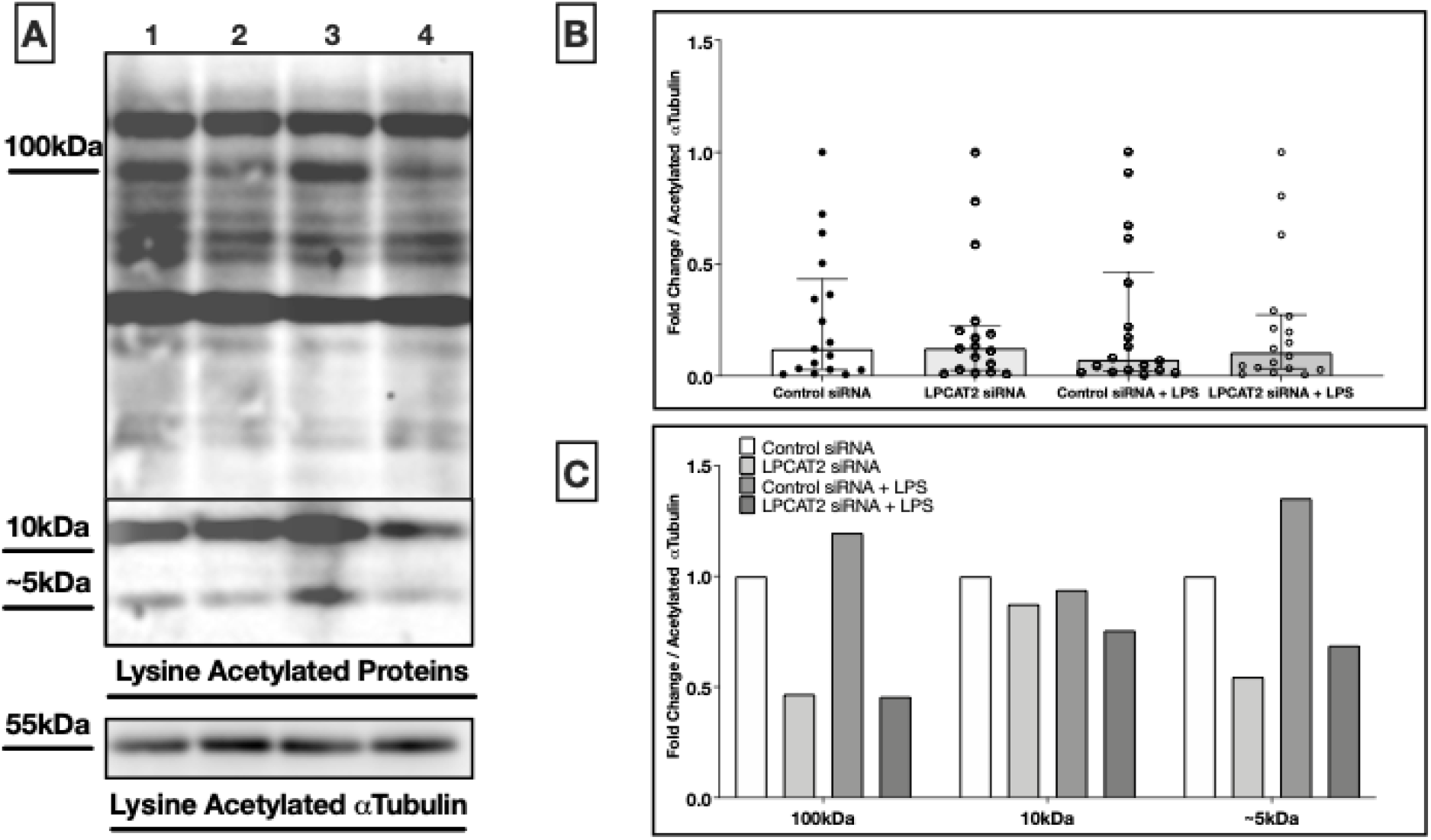
Analysis of Pan-Lysine Acetylation in Lipopolysaccharide-Stimulated RAW264.7 Cells. **(A)** – Western blot image of lysine acetylated proteins. Acetylated Alpha Tubulin was used as a positive control. Lane 1 - Control siRNA, Lane 2 - LPCAT2 siRNA, Lane 3 - Control siRNA + LPS, Lane 4 - LPCAT2 siRNA + LPS. **(B)** – Fold change in optical density of lysine acetylated proteins normalised to acetylated αTubulin. Bars represent median and error bars represent interquartile range. **(C)** – Fold change in optical density of bands at 100kDa, 10kDa, and ∼5kDa normalised to acetylated αTubulin and control siRNA.

### 4.4 In Silico Prediction of Lysine Acetylation of Lipopolysaccharide Receptors

Due to the presence of acetylated lysine which can be induced by LPS and decreased by silencing LPCAT2 at 100kDa, the protein sequences of mouse TLR4 which is about 100kDa in size and its co-LPS-receptors; MD2 and CD14 were analysed for the possible presence of lysine residues that can be acetylated. Two software were used for this analysis; GPS-PAIL version 2.0 and ASEB. Table 1 shows that GPS-PAIL version 2.0 software predicted that TLR4 is acetylated on the lysine residue at position 817 by CREBBP, but CD14 and MD2 did not show any possibility for lysine acetylation. RelA was used as a positive control, as it is already experimentally proven to undergo lysine acetylation [26].

**Table 1:** *In Silico* Prediction of Lysine Acetylation in lipopolysaccharide Receptors Using GPS-PAIL Version 2.0.

| Protein | Position | Peptide | Score | Cut-off | Acetyltransferase |
| --- | --- | --- | --- | --- | --- |
| CD14 | NA | NA | NA | NA | NA |
| MD2 | NA | NA | NA | NA | NA |
| TLR4 | 817 | KNALLDGKASNPEQ | 2.09 | 1.79 | CREBBP |
| RelA | 310 | KRTYETFKSIMKKS | 2.32 | 1.79 | CREBBP |
| RelA | 122 | NLGIQCVKKRDLEQ | 2.08 | 1.69 | KAT2B |

NA indicates that no sites of lysine acetylation was detected. Position indicates the position of the lysine with predicted acetylation. The acetyltransferases were also predicted based on known information about binding sites of these enzymes. The score indicates the probability of acetylation. The higher the score, the higher the chance of acetylation. The cut-off is the number under the threshold. This analysis was carried out using a high threshold. Red letters signify the supposed position of lysine acetylation.

Further analysis of mouse TLR4 protein sequence using ASEB revealed the possibiility of more than 1 lysine residue undergoing lysine acetylation. In Table 2, the lysine residue at position 817 was predicted to undergo lysine acetylation by CREBBP along with lysine residues at positions 367 and 503. The lysine residue at position 152 was predicted to undergo lysine acetylation by KAT2A and or KAT2B.

**Table 2:**
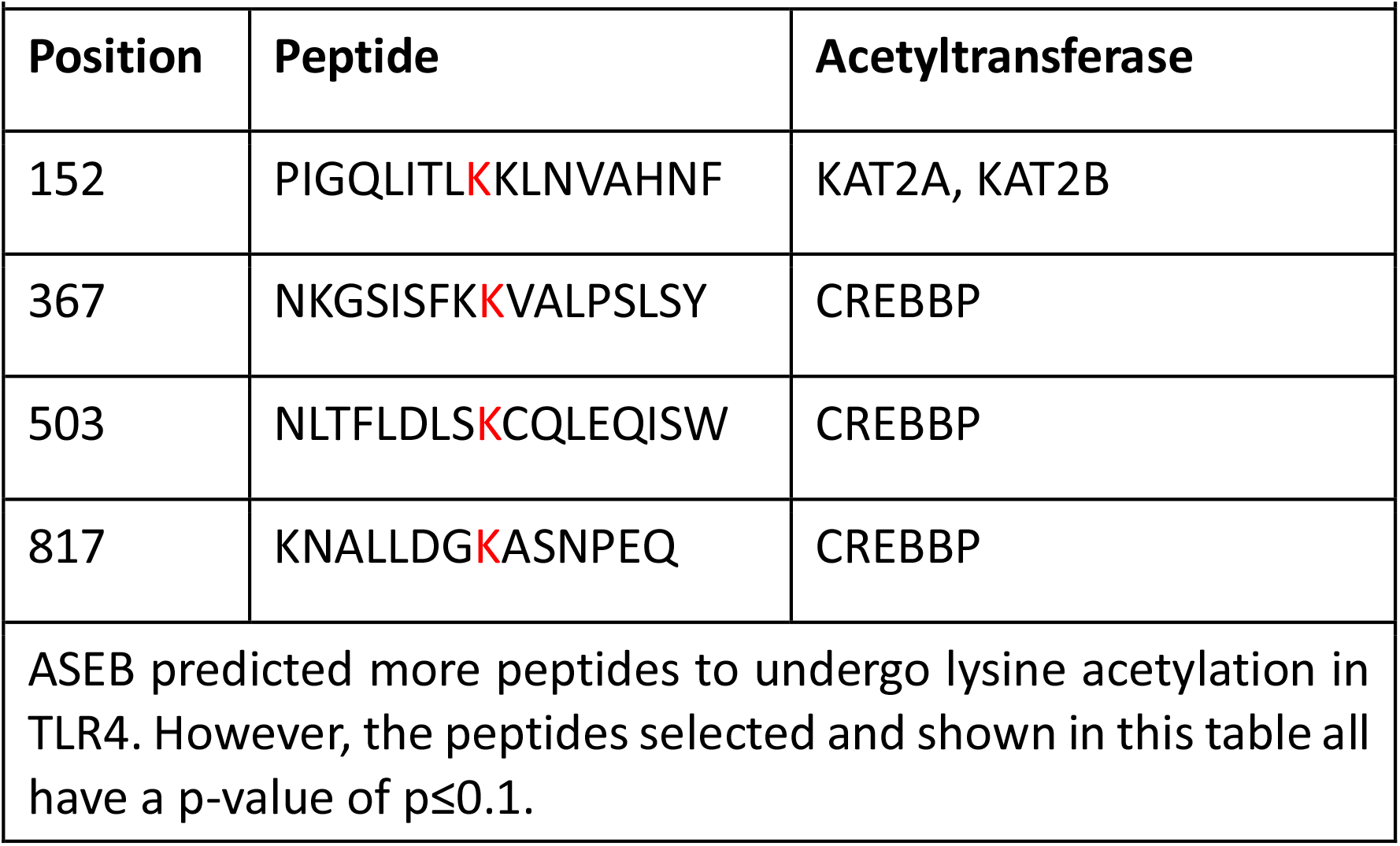
*In Silico* Prediction of Lysine Acetylation of TLR4 ASEB.

To understand the properties of the predicted peptides and predict the potential roles of the acetylated lysine residues, analysis of each peptide predicted to contain acetylated lysine residue was carried. As shown in Table 3, all peptides were globular, suggesting that the peptides are situated either on the extracellular or the intracellular end of TLR4, but not the transmembrane region. The high aliphatic index values also predict that these peptides are thermostable.

**Table 3:**
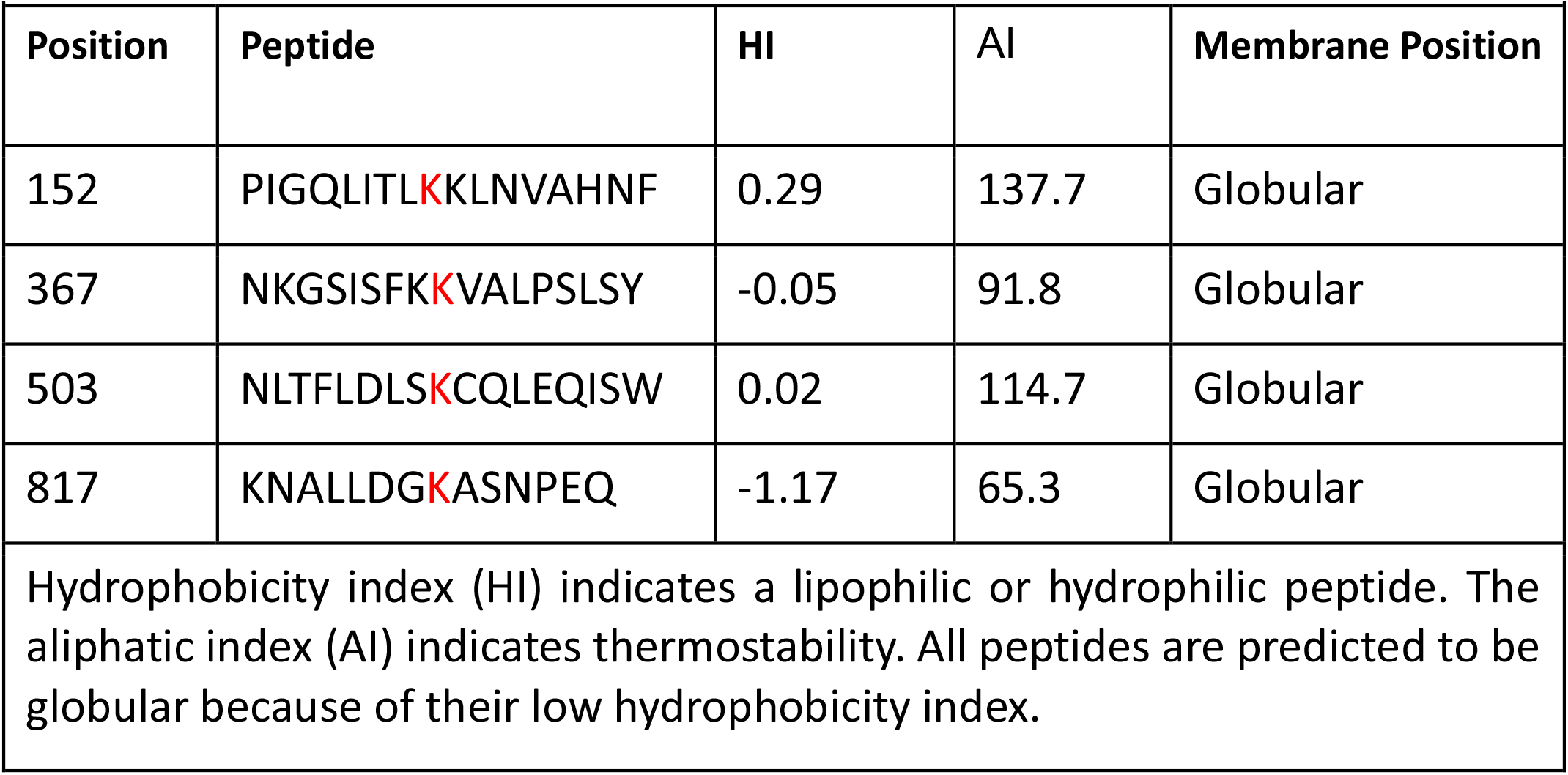
Properties of Predicted Lysine Acetylated Peptides in TLR4.

### 4.5 LPCAT2 Influences the Lysine Acetylation of Toll-like Receptor 4

Further analysis to confirm the presence of lysine acetylation in TLR4 was carried out by immunoprecipitating TLR4 and blotting for acetylated lysine residues. Figure 4A shows that at 100kDa, TLR4 was detected in both blots using TLR4 antibodies and acetylated lysine antibodies after immunoprecipitation of TLR4. In Figure 4B, the optical densities of TLR4 in both TLR4 blots and acetylated lysine blots were both within the range of 3000 to 4500. Treatment of RAW264.7 cells with LPS did not show any difference.

**Figure 4:**
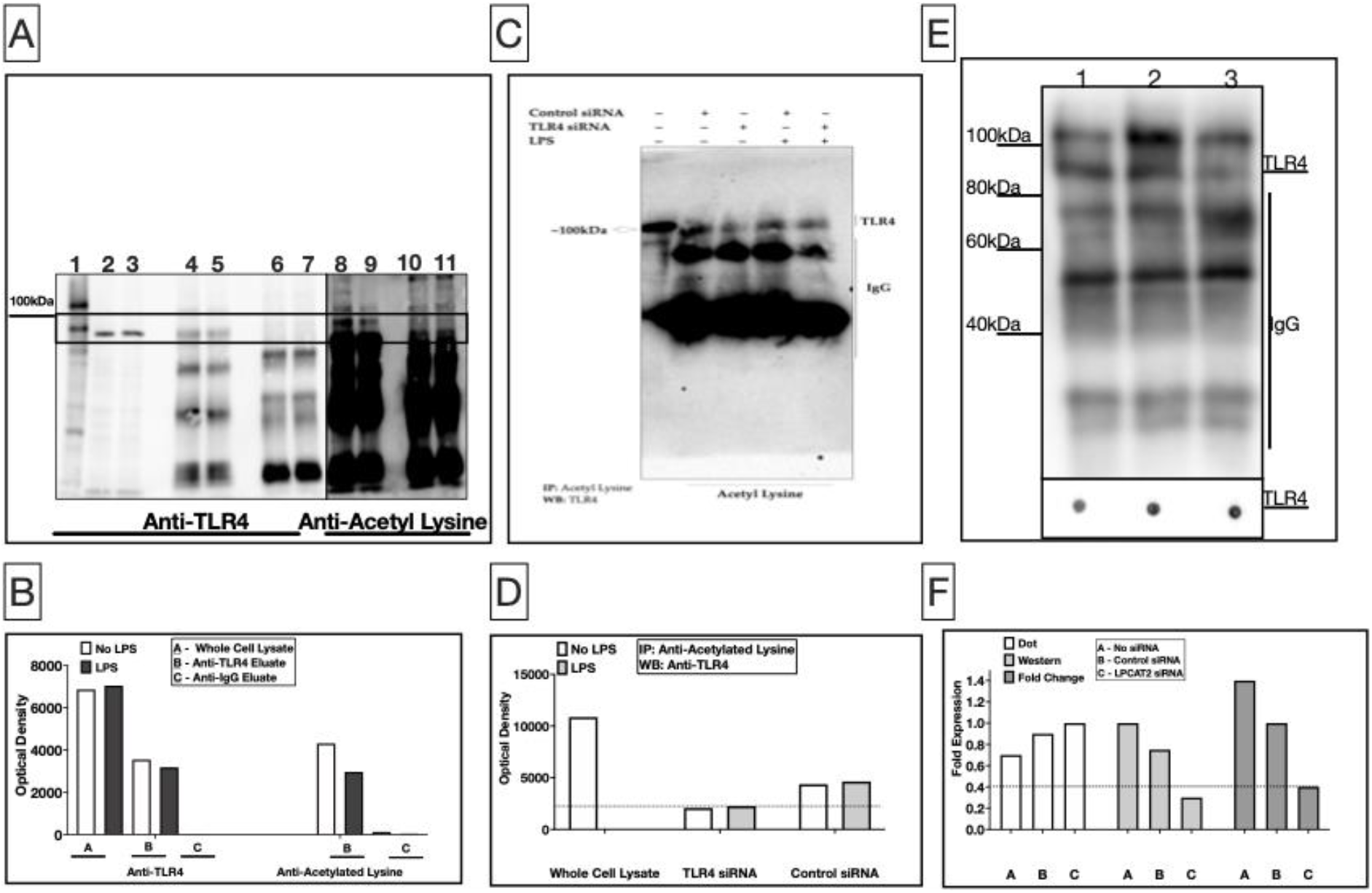
**A–** Acetylated Lysine Detected on Precipitated TLR4. Lane 1-biotinylated protein ladder; Lanes 2 & 3 - TLR4 blots of whole cell lysates; Lanes 4 & 5 -TLR4 blots of TLR4 eluate; Lanes 6 & 7 - TLR4 blots of IgG eluate; Lanes 8 & 9 -acetylated lysine blots of TLR4 eluate; Lanes 10 & 11 - acetylated lysine blots of IgG eluate. Lanes 2, 4, 6, 8, and 10 are samples from RAW264.7 cells not treated with LPS, whereas lanes 3, 5, 7, 9, and 11 are samples from RAW264.7 cells treated with 100ng/ml LPS. **B–** Optical densities of protein bands at 100kDa. Anti-IgG elution was used as isotype control. **C–** TLR4 blot (WB) of acetylated lysine precipitates (IP). The first lane is a blot of lysate of RAW264.7 cells not transfected with siRNA or treated with LPS. The other lanes are blots of acetylated lysine eluates. - indicates absence, and + indicates presence. **D–** Optical densities of protein bands at 100kDa.**E–** Acetylated lysine blots (WB) of TLR4 precipitates (IP). Lane 1 - No siRNA, Lane 2 - Control siRNA, Lane 3 - LPCAT2 siRNA. TLR4 dot blots of uniform amounts of proteins from TLR4 eluates were used as controls of protein amounts. **F–** Optical density of protein bands at ∼90kDa. Fold change refers to values normalised to dot blot of TLR4 in eluates and to control siRNA.

In an attempt to ensure that the protein being detected is TLR4, we silenced the expression of TLR4 in RAW246.7 cells and repeated the analysis. In this case, acetylated lysine proteins were immunoprecipitated and eluates were analysed for the presence of TLR4. The postulation was that if the detected protein was not TLR4, then it will not decrease when TLR4 is silenced and it will not be detected by TLR4 antibody. In Figure 4C, we see again that TLR4 was detected from eluate of lysine acetylated proteins by TLR4 antibody. Figure 4D shows again that knockdown of TLR4 reduced the presence of lysine acetylation on TLR4 by ≥50%. This is further evidence of the presence of lysine acetylation on TLR4. TLR4 expression was also reduced in cells not treated with LPS but transfected with TLR4 siRNA (Poloamina, 2021).

As the initial study was to understand the role of LPCAT2 on lysine acetylation, we sought to determine the influence of LPCAT2 on the detected lysine acetylation on TLR4. Figure 4E, shows a TLR4 blot of acetylated lysine eluates from RAW264.7 cells. The optical densities in Figure 4F shows ≥50% decrease in acetylated TLR4 lysine after knockdown of LPCAT2.

### 4.6 LPCAT2 Regulates the Expression of Interferon Beta and Interferon-Inducible Protein 10 (IP10)

A recent publication has shown that LPCAT2 can regulate the expression of pro-inflammatory cytokines; TNFα and IL6 (Abate et al, 2020), however, interferon-beta (IFNβ) and interferon-inducible protein 10 (IP10) which are dependent on TLR4-Trif signalling pathway, were not analysed. Figure 5A shows that knockdown of LPCAT2 results in a significant decrease (5.38 ± 0.69, p = 0.00025) in the gene expression of IFNβ in cells stimulated with 100ng/ml lipopolysaccharide. IP10 which can be also be induced by IFNβ, was analysed. Figure 5B shows that knockdown of LPCAT2 significantly decreased the gene expression (168.7 ± 32.27, p = 0.021). In non-treated cells, knockdown of LPCAT2 did not effect a significant change in IFNβ gene (p = 0.46), and IP10 gene (p = 0.97) expression.

**Figure 5:**
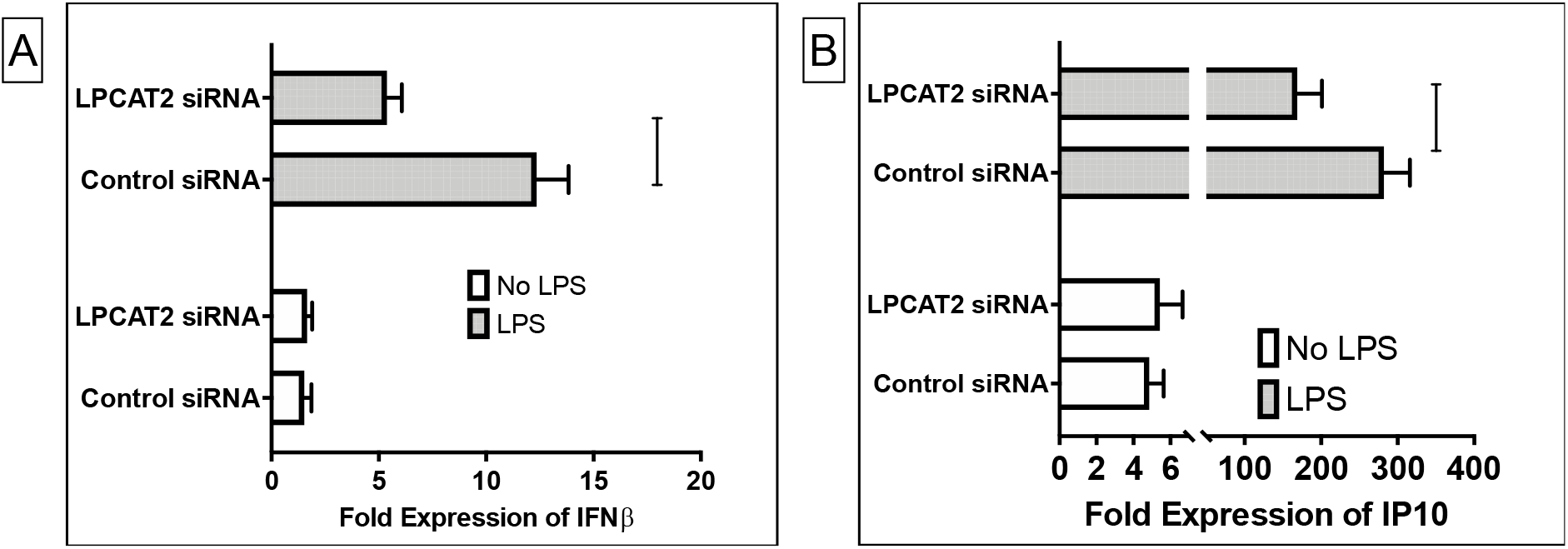
Knockdown of LPCAT2 affects the gene expression of Interferon beta, IFNβ **(A)** and interferon-inducible protein 10, IP10 **(B)** in RAW264.7 cells stimulated with TLR4 ligand (100ng/ml of E. coli O111:B4 lipopolysaccharide). Non-treated RAW264.7 cells did not show any significant difference. *p≤0.05. Data represents the mean of at least 3 independent experiments (n≥3) ± standard error.

## 5 Discussions

LPCAT2 is commonly known as a lipid-modifying enzyme. However, there is scientific evidence that LPCAT1, which is similar in structure and function to LPCAT2, palmitoylates histone 4 [22]. This implies that LPCAT2 may modify proteins. Moreover, LPCAT2 co-localises with TLR4 and modifies its subcellular localisation and hence its function, through an unknown mechanism [5]. Addition of acyl groups to proteins can regulate its subcellular localisation and or function [27]. Using both computational and biochemical analysis, our results suggest that TLR4-LPCAT2 co-localising may result in the lysine acetylation of TLR4 which may influence its subcellular localisation.

### 5.1 LPCAT2 Gene and Protein Sequence is homologous to other lysine acetyltransferases

50% to 70% homology is required for conservation of enzyme functions [28]. Figure 2 shows that LPCAT2 shares at least 65% similarity with KAT2A and KAT2B, which are established lysine acetyltransferases. Moreover, analysis of their gene sequences showed that LPCAT2, LPCAT1, KAT2A, and KAT2B evolved from the same superfamily. The results from this computational analysis lays a foundation for further scientific experiments that will be required to classify LPCAT2 as a lysine acetyltransferase.

### 5.2 LPCAT2 Influences Lysine Acetylation Detected on TLR4 Protein

The housekeeping genes– ATP5B and GAPDH and the total RNA and protein amount did not undergo any significant changes when RAW264.7 cells were transfected with siRNA (Figure 1). This eliminates the possibility that the observed gene expression or protein expression is as a result of overall changes in the cell.

Palmitoylation can regulate the subcellular localisation and signalling of TLR2. It is also present on other TLRs like TLR5 and TLR10, but not on TLR4 [29]. As TLR4 was not found amongst palmitoylated proteins, we explored the next possibility which is acetylation. LPCAT2 has both acyltransferase and acetyltransferase [6], however, unlike LPCAT1 there is no previously published evidence that suggests that LPCAT2 can carry out its enzymatic activities on proteins. Our results show that when LPCAT2 is knocked down in RAW264.7 cells, the band intensities of some lysine acetylated proteins reduces especially at 100kDa and 5kDa (Figure 3). Except TLR4, there are many other proteins with an approximate mass of 100kDa. On the UniProt database [30], there are about 3500 proteins between 90kDa and 110kDa. Only about 25 of these proteins have published evidence of lipidation. Therefore, a more TLR4-specific study is needed to confirm its lysine acetylation. Computational analysis to predict TLR4 lysine acetylation suggests that TLR4 may undergo lysine acetylation on lysines 152, 367, 503, and 817 cataysed by KAT2A, KAT2B, or CREBBP (Table 1 & Table 2). Biochemical analysis shows the presence of lysine acetylation on TLR4 (Figure 4), however, further experiments will be required to know the specific lysine residues that are acetylated on TLR4 and their role in regulating TLR4 function. In Table 3, we predicted that the TLR4 peptides that have acetylated lysine residues are hydrophilic and globular. Although this is not confirmatory, it suggests that lysine acetylation may be occuring on the extracellular domain of TLR4 where it binds to its ligands, or the intracellular domain of TLR4 where it binds to other proteins and transmits signals. Indeed, a thesis suggested that TLR4 lysine acetylation influences its interaction with its adaptor proteins MyD88, TRAM, and TRIF; which eventually affects the production of inflammatory cytokines [31]. We have previously shown that knockdown of LPCAT2 reduces MyD88-dependent cytokines after LPS-TLR4 interaction [5] Likewise, Figure 5 shows that TRIF-dependent cytokines are influenced by knockdown of LPCAT2. TLR4 protein or gene expression is not affected by silencing LPCAT2 (data not shown), however, Figure 4 suggests for the first time that LPCAT2 may influence lysine acetylation detected on TLR4.

In conclusion, the results from this study suggest that TLR4 undergoes lysine acetylation and LPCAT2 influences the detected lysine acetylation. This lays a foundation for further research on the role of lysine acetylation on TLR4, and characterisation of LPCAT2 as a protein acetyltransferase.

